# A “Dirty” Footprint: Anthropogenic Soils Promote Biodiversity in Amazonian Rainforests

**DOI:** 10.1101/552364

**Authors:** Wilian C. Demetrio, Ana C. Conrado, Agno N.S. Acioli, Alexandre Casadei Ferreira, Marie L.C. Bartz, Samuel W. James, Elodie da Silva, Lilianne S. Maia, Gilvan C. Martins, Rodrigo S. Macedo, David W.G. Stanton, Patrick Lavelle, Elena Velasquez, Anne Zangerlé, Rafaella Barbosa, Sandra Celia Tapia-Coral, Aleksander W. Muniz, Alessandra Santos, Talita Ferreira, Rodrigo F. Segalla, Thibaud Decaëns, Herlon S. Nadolny, Clara P. Peña-Venegas, Cláudia M.B.F. Maia, Amarildo Pasini, André F. Mota, Paulo S. Taube Júnior, Telma A.C. Silva, Lilian Rebellato, Raimundo C. de Oliveira Júnior, Eduardo G. Neves, Helena P. Lima, Rodrigo M. Feitosa, Pablo Vidal Torrado, Doyle McKey, Charles R. Clement, Myrtle P. Shock, Wenceslau G. Teixeira, Antônio Carlos V. Motta, Vander F. Melo, Jefferson Dieckow, Marilice C. Garrastazu, Leda S. Chubatsu, TPI Network, Peter Kille, George G. Brown, Luís Cunha

## Abstract

Amazonian rainforests once thought to hold an innate pristine wilderness, are increasingly known to have been densely inhabited by populations showing a diverse and complex cultural background prior to European arrival. To what extent these societies impacted their landscape is unclear. Amazonian Dark Earths (ADEs) are fertile soils found throughout the Amazon Basin, created by pre-Columbian societies as a result of more sedentary habits. Much is known of the chemistry of these soils, yet their zoology, have been neglected. Hence, we characterised soil macroinvertebrate communities and activity in these soils at nine archaeological sites in three Amazonian regions. We found 667 morphospecies and a tenacious pre-Columbian footprint, with 43% of species found exclusively in ADEs. The soil biological activity is higher in the ADEs when compared to adjacent reference soils, and it is associated with higher biomass and richness of organisms known to engineer the ecosystem. We show that these habits have a unique pool of species, however, the contemporary land-use in ADEs drives nutrient decay and threats biodiversity. These findings support the idea that Humans have built and sustained a contrasting high fertile system that persisted until our days and irreversibly altered the biodiversity patterns in Amazonia.

---

The Amazon basin contains the largest continuous and relatively well-preserved tract of tropical forest on the planet. Although deforestation rates in Amazonia have been showing a generally decreasing trend over the last decade, human activities in the region were still responsible for losses of 7,900 km^2^ of its natural vegetation in 2018 alone^1^. Many forested areas have become highly fragmented, and may be reaching tipping points where biodiversity and ecosystem functions may be dramatically affected^2, 3^, potentially leading to cascading effects that impact ecosystem services over a much larger area^4, 5^.

Humans have modified Amazonian biodiversity patterns over millennia, and Amerindians created areas with high concentrations of useful trees and hyperdominance of some species, often associated with archaeological sites^6^ (Fig. 1a). Furthermore, occupations of some indigenous societies’, beginning at least 6,500 years ago, created fertile anthropogenic soils, locally called “*Terra Preta de Índio*” (TPI) or Amazonian Dark Earths – ADEs^7–9^ (Fig. 1b). The ADEs may occupy up to 3% of the surface area of Amazonia^7^, and appear to be more common along major rivers (Fig. 1a), but are also abundant in interfluvial areas^9^. ADE sites tend to have high soil P, Ca and pyrogenic C contents^10–12^, and particular communities of plants and soil microorganisms^13, 14^, but up to now, soil animal communities in these historic anthropogenic soils were not previously known.

**Figure 1.**
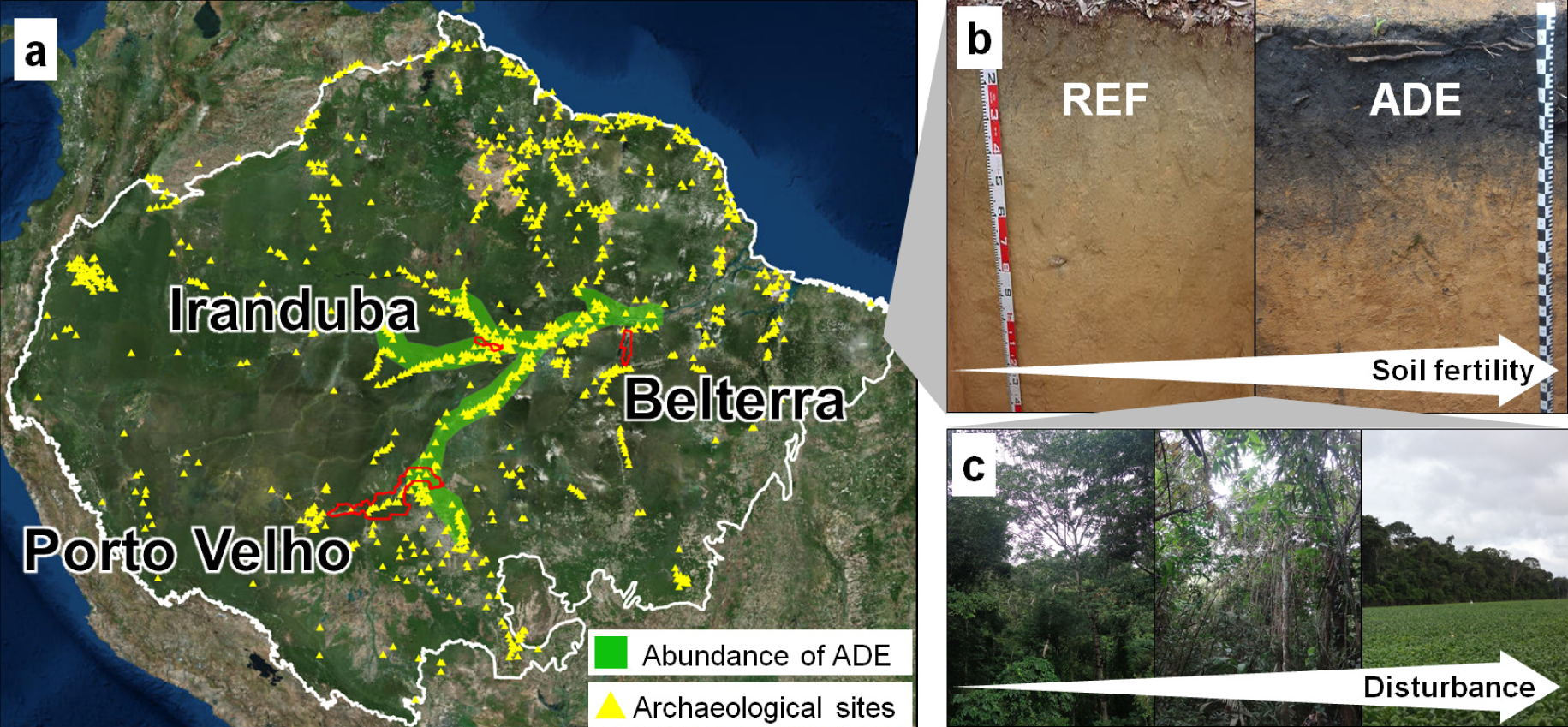
Sampling strategy to assess soil fauna and soil fertility in Central (Iranduba), Southwestern (Porto Velho) and Lower (Belterra) Amazon. (**a**) Boundary of Amazon Basin (white line), boundaries of municipalities where samples were taken (red lines), archaeological sites (yellow triangles), and areas with high concentration of Amazonian Dark Earths (ADE, shaded in green) at archaeological sites. Archaeological and ADE sites modified from Clement et al.^9^ Amazonia map background: Esri, DigitalGlobe, GeoEye, Earthstar Geographics, CNES/Airbus DS, USDA, USGS, AEX, Getmapping, Aerogrid, IGN, IGP, swisstopo, and the GIS User Community. (**b**) Soil profiles of analytically paired ADE and nearby reference (REF) soils; Photos G.C. Martins, R. Macedo. (**c**) Land use systems (LUS) sampled in each region, consisting in an intensification/disturbance gradient including old secondary rainforest (>20 yrs. undisturbed), young secondary forest (<20 yrs. old), and recent agricultural systems (pasture, soybean, maize); Photos G.C. Martins, M. Bartz.

Soil macroinvertebrates represent as much as 25% of all known described species^15^, and are a huge source of biodiversity that may easily surpass 1 million species^16^. However, soil animal communities have been little studied in megadiverse regions, such as the Amazonian rainforest^17, 18^, and these habitats may be home to thousands of species^19, 20^, particularly smaller invertebrates such as nematodes and mites^21, 22^, but also of macroinvertebrates.

Hence, the aim of this study was to assess soil invertebrate macrofauna communities and their activity in ADEs at nine archaeological sites and adjacent reference soils (REF) under three land-use systems (LUS: old and young secondary forest and recent agricultural/pastoral systems), in order to evaluate anthropic effects on Amazonian soil biodiversity. We predicted that (1) soil biodiversity composition and soil enrichment in anthropogenic soils would reflect a pre-Colombian footprint but also, that (2) animal richness, biomass, activity, and nutrient contents in these soils would be determined by present-day land-use.

## Results

### ADEs are distinct soils with distinctive macroinvertebrate communities

The ADEs at all the sites had higher soil pH and were enriched in Ca, Mg, P and total C compared to REF soils within each LUS (Fig. 2), following trends typically observed in ADE sites throughout Amazonia^12, 23^. Significantly lower amounts of exchangeable Al were also found in the ADEs (Supplementary Table 1). Soil texture at the sites was similar in both ADE and REF soils (Supplementary Table 1), so the enrichment was not due to differential clay contents, but the result of ancient anthropogenic activities^23, 24^. Some differences in soil fertility among land-use systems were also observed (Supplementary Table 1), where plots under agricultural or pastoral use (AS) had higher K contents (due to fertilization) than old forest (OF) and lower N contents, probably due to soil erosion processes, denitrification, and leaching^25, 26^.

**Figure 2.**
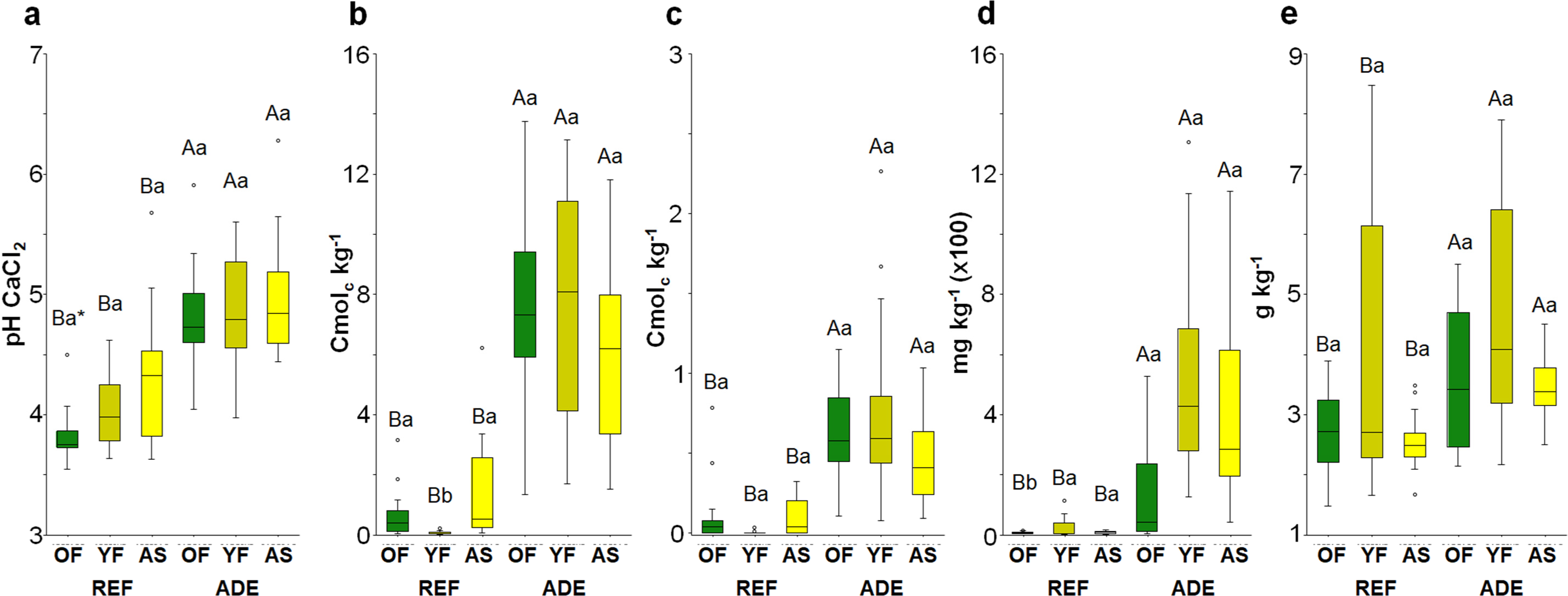
Soil chemical properties at the collection sites in Amazonia: (**a**) pH, (**b**) exchangeable Ca, (**c**) exchangeable Mg, (**d**) available P, and (**e**) total carbon in the topsoil layer (0-30 cm depth; mean values for the three regions) showing differences due to soil type (REF vs. ADE soils) in each of the land-use systems (OF, YF, AS). *Different upper-case letters indicate significant differences (*P* < 0.05) among soil categories within each land-use system, while different lower-case letters indicate significant differences among land use systems within the same soil type. ADE: Amazonian Dark Earth; REF: reference soils; OF: old forests; YF: young forests; AS: agricultural systems. Values shown are median (black line), 1^st^ and 3^rd^ quartiles (box) and max/min observations (upper and lower lines) and the outliers (small open circles), when present.

We collected 9,380 macroinvertebrates in soil monoliths, of 667 different morphospecies, belonging to 24 higher taxa (Fig. 3a; Supplementary Table 2). Ants (Formicidae) were the most diverse group collected (154 spp.), followed by spiders (86 spp.), beetles (78 spp.), millipedes (53 spp.), true bugs (42 spp.), termites (37 spp.), cockroaches (34 spp.), and earthworms (32 spp.) (Supplementary Table 2). The number of singleton species (one individual in the total sample of 9,380) was very high (328 spp.), representing around 49% of the total macroinvertebrate richness (Supplementary Table 3).

**Figure 3.**
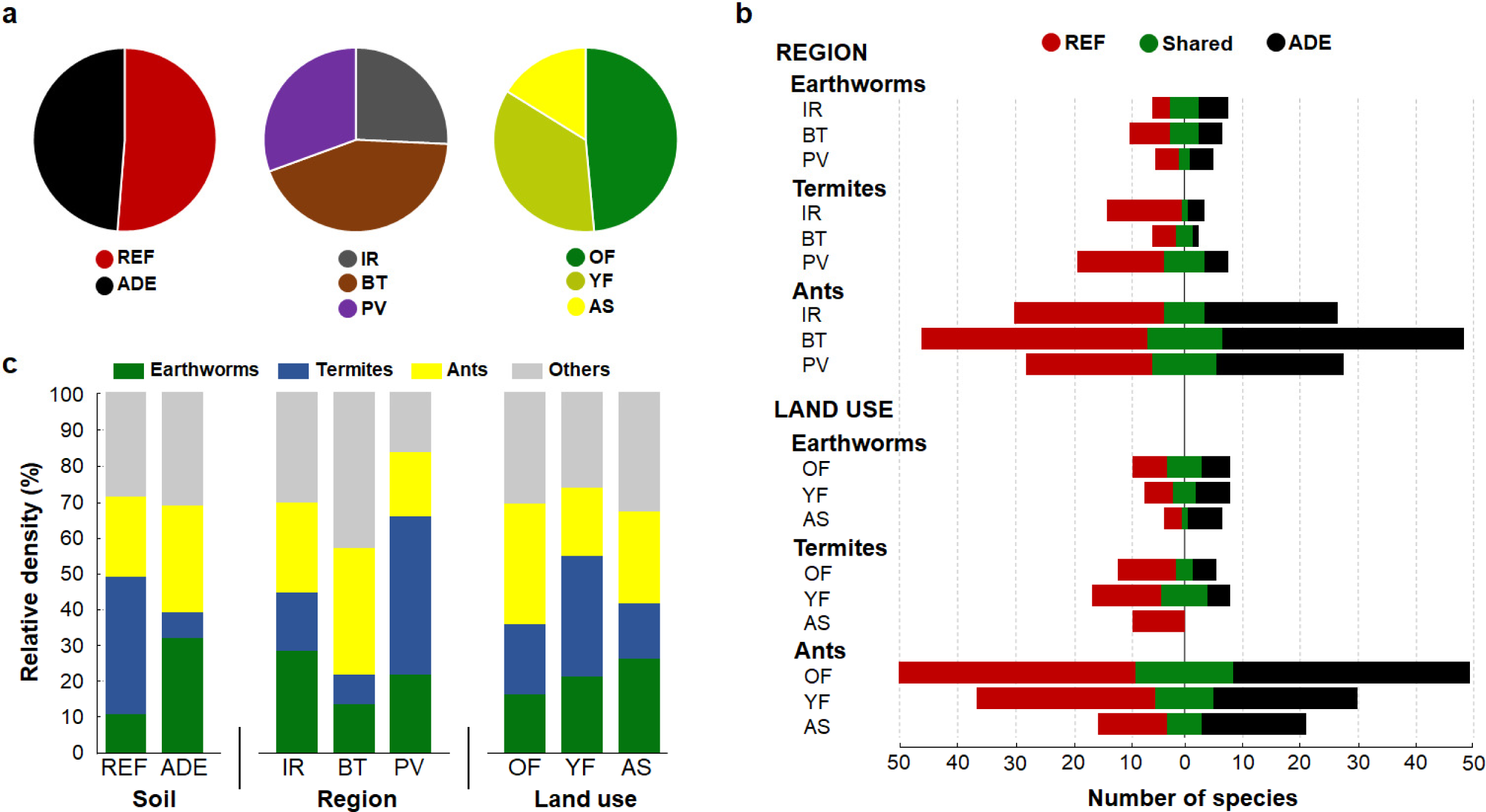
Morphospecies diversity and abundance patterns in soil communities at collection sites in Amazonia: (**a**) Distribution of unique morphospecies (including singletons) of all macroinvertebrates according to soil type (ADE, REF), region (IR, BT, PV) and land use systems (OF, YF, AS). (**b**) Numbers of morphospecies of earthworms, termites and ants observed in both soil categories (green bars) or uniquely in ADE (black bars) or in REF (red bars) soils, in the different regions and land use systems across regions. (**c**) Relative density (%) of earthworms, termites, ants and other soil macroinvertebrates (sum of all other taxa) found in the different soil categories (ADE vs. REF), regions, and land use systems; ADE: Amazonian Dark Earth; REF: reference soils; IR: Iranduba; BT: Belterra; PV: Porto Velho; OF: old forests; YF: young forests; AS: agricultural systems.

Similar numbers of species were found in ADEs (382 spp.) and REF (399 spp.) soils. The proportion of unique morphospecies was high in both soils: 48.5% in ADEs and 51.5% in REF soils (Fig. 3a; Supplementary Fig.1), particularly for ants (75 spp. ADE, 70 spp. REF) and earthworms (22 spp. ADE, 20 spp. REF) (Fig. 3b; Supplementary Figs 2-4). Termites had a high number of unique species in REF soils (21 spp.; see Fig. 3b). These trends for ants, earthworms and termites remained similar even after singleton species were removed. Centipede and Opiliones richness was also high in REF soils (14 and 14 spp., respectively), while millipede and snail richness (37 spp. and12 spp., respectively) was high in ADEs (Supplementary Table 2), possibly due to the higher soil Ca levels^27^. The high number of species unique to each soil (Fig. 3a) was reflected in high β-diversity values and species turnover, ranging from 67-79% for all of the soil macroinvertebrates (Supplementary Table 6). Furthermore, among the ecosystem engineers collected, we found an important number of species new to science (>20 earthworm species, >20 termite species and >30 ant species) that still need to be described.

ADEs were home to 95 rare (doubleton and rare individuals) and to 18 non-rare or abundant macroinvertebrate morphospecies not found in REF soils (Supplementary Table 3). Interestingly, within the non-rare/abundant taxa, 19 species (mainly ant and earthworm species) had greater abundance of individuals in ADEs, while 13 species (mainly ant species) were more prevalent in REF soils (Supplementary Table 3).

Estimated richness for total macroinvertebrates, ants and earthworms (Fig. 4a,b,d) was not different between REF and ADE soils but for termites was two-times higher in REF soils (Fig. 4c). These results were confirmed with the more intensive sampling effort performed for ants, termites, and earthworms (Supplementary Fig. 5). The monolith samples’ collected around 65-75% of the estimated richness of total soil macroinvertebrates and ants in both soil types and of termites in REF soils (Supplementary Fig. 6a,b,c). Earthworm richness in both soil categories and termite species in ADEs were relatively well sampled by the monoliths, which collected 70-80% of the estimated total diversity (Supplementary Fig. 6c,d). The use of complementary sampling methods increased the number of collected species for ants in both soils and for termites in REF soils (Supplementary Fig. 5a,b), revealing an important un-sampled species pool of these soil engineers (particularly of ants) in the forests of each region, especially in REF soils.

**Figure 4.**
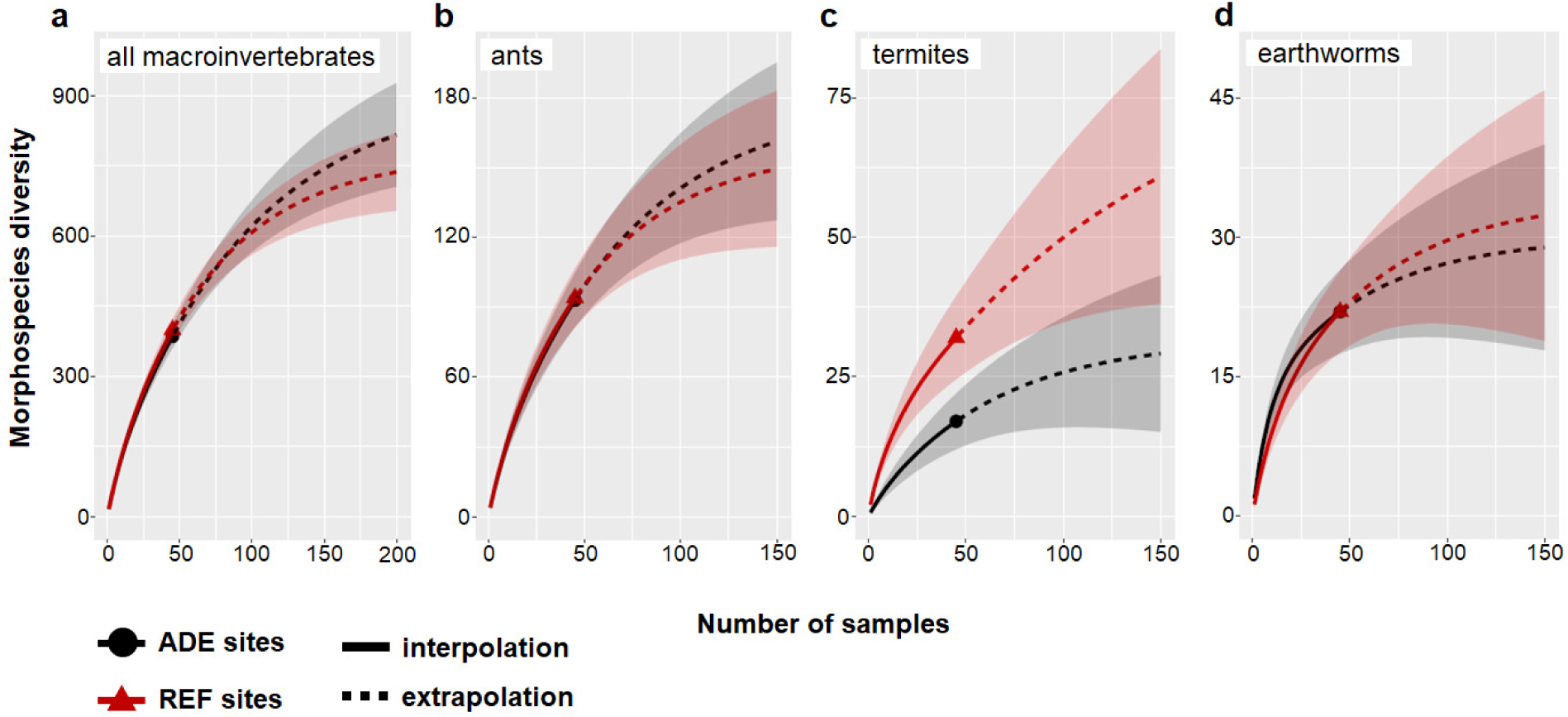
Morphospecies rarefaction and extrapolation curves showing how morphospecies quantities increase in both ADE and REF soils depending on sampling intensity (number of samples) for: (**a**) all soil macroinvertebrates, (**b**) ants, (**c**) termites and (**d**) earthworms. Data correspond to invertebrates collected in soil monoliths from all sites and land use systems. Dark grey and red areas represent 95% confidence intervals. ADE: Amazonian Dark Earth; REF: Reference soil.

Land-use effects on species turnover rates were slightly higher for all soil macroinvertebrates (0.79 and 0.74 within REF and ADEs, respectively) than for soil type comparisons (0.70, 0.67 and 0.71 for OF, YF and AS, respectively), indicating that species turnover was more closely related to LUS than to soils (Supplementary Table 6). Similar results were observed for earthworms, with much higher turnover rates (0.84 and 0.62 within REF and ADEs, respectively) due to LUS than due to soil, particularly in OF and YF. Conversely, soil type had a greater impact on ant and termite species turnovers than land-use (0.78 for ants and 0.72 for termites in OF). The species turnover among regions was also very high, mainly for overall macroinvertebrates and earthworms in AS (Supplementary Table 7).

### Ecosystem engineers dominate the soil fauna communities

Ecosystem engineers (termites, ants and earthworms)^31^ represented on average 72% and 69% of the soil macroinvertebrate individuals in ADE and REF soils, respectively (Fig. 3c). The proportion of ecosystem engineers was significantly higher in PV than IR and BT, mainly due to the higher proportion of termites in PV (Fig. 3c). Ecosystem engineers represented 62 to 75% of total invertebrate biomass in the different LUS and soil categories, and was not significantly different between ADE and REF soils (Supplementary Table 4). Termite populations were significantly higher in REF soils with populations over 1000 individuals m^−2^, while earthworms, ants, and other invertebrates were proportionally more prevalent in ADE (Fig. 3c; Supplementary Table 4). Ants were proportionally more abundant at BT, and termites in IR and PV (Fig. 3c). In biomass, earthworms represented from 44% (AS, REF) to 92% (AS, ADE) of the total macroinvertebrate biomass, and their abundance and biomass were significantly higher in ADE (particularly in YF and AS) than in REF soils (Supplementary Table 4). No other soil animal represented more than 35% of the biomass in any given soil type or LUS.

### Modern land use erodes soil biodiversity

A total of 349, 278, and 152 morphospecies of macroinvertebrates were found in OF, YF and AS, respectively, of which 249, 181, and 83 species were unique to the respective LUS (Fig. 3a). Removing singleton species, morphospecies richness was 137 (OF), 98 (YF) and 47 (AS) in ADE, and 122 (OF), 102 (YF) and 54 (AS) in REF soils. Hence, richness was 56% and 46% lower in modern AS compared with OF and YF, respectively. This trend was also observed for most of the groups of soil animals, and was particularly marked (>60% decrease in spp. richness) for opilionids, centipedes, isopods and cockroaches in both REF and ADEs, and for earthworms in REF and termites in ADEs (Supplementary Table 2). Species richness decreases in AS compared to OF were slightly (but not significantly) higher for ADE (66%) and REF (56%) soils.

### Soil biota influence ADE soil structure

Soil macromorphology revealed a significantly higher proportion of fauna-produced aggregates (Fig. 5) in ADE soils compared with REF soils, and likewise, in the same LUS, a lower proportion of non-aggregated soil (Supplementary Table 5) in ADEs than REF soils, implying important changes in soil structure in ADEs. Fauna-produced aggregates were also more abundant in OF compared to YF and AS systems (Fig. 5), which tended to have higher proportions of loose, non-aggregated soil and physical aggregates (Supplementary Table 5). The proportions of other aggregate fractions were not affected by soil type and LUS (Supplementary Table 5).

**Figure 5.**
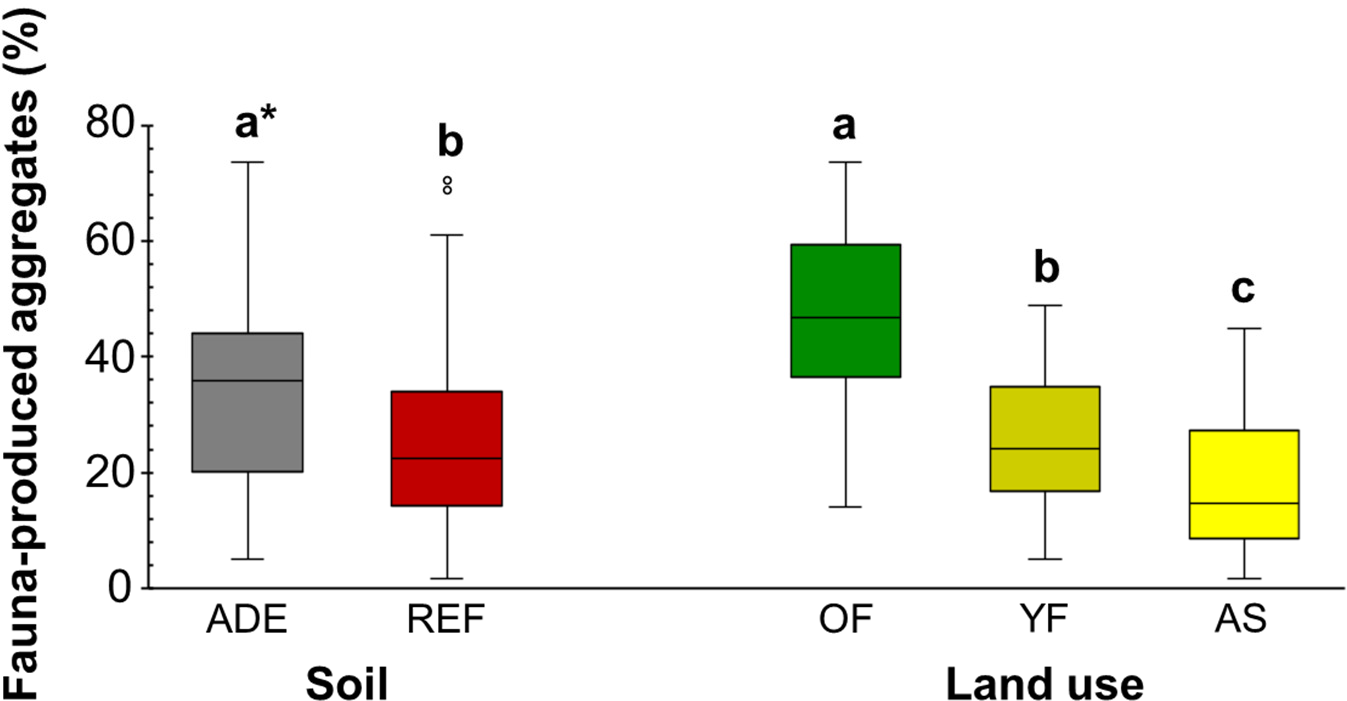
Proportion of fauna-produced aggregates in top-soil. (0-10 cm layer) in two different Amazonian soils (REF: non-anthropogenic reference soils; ADE: from Amazonian Dark Earth) and three different land use systems (OF: old forests, YF: young forests, AS: agricultural systems). Values shown are median (black line), 1^st^ and 3^rd^ quartiles (box) and max/min observations (upper and lower lines) and the outliers (small open circles), when present. *Different letters indicate significant differences (*P* < 0.05) within soil or land use comparisons.

Multivariate analysis (PCA) confirmed the importance of soil fertility associated with ADE (nutrient contents aligned with x-axis) and REF soils as a regulator mainly of earthworm and termite abundance, and land use disturbance or intensification (LUS aligned with y-axis) as a regulator of ant and overall soil fauna abundance and biodiversity (Fig. 6).

**Figure 6.**
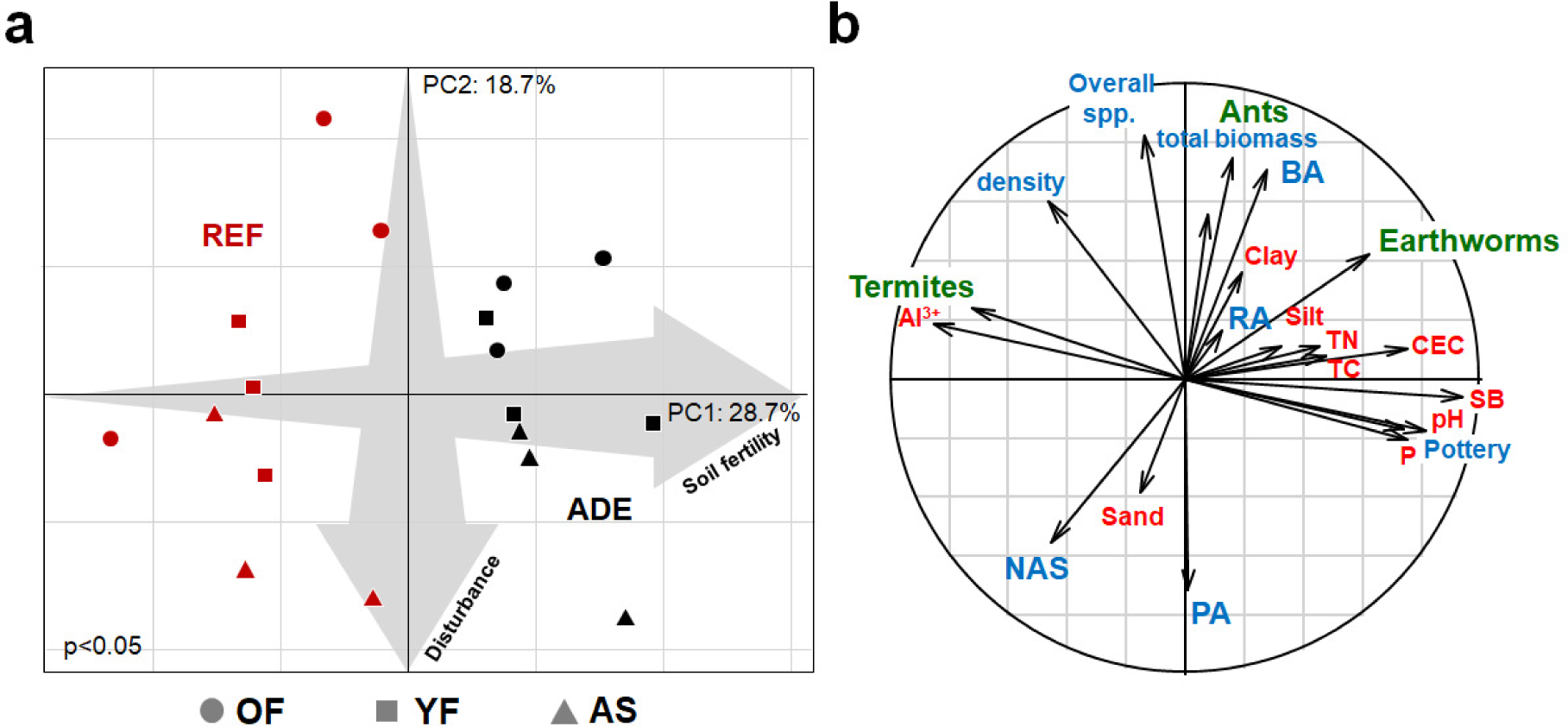
Principal Component Analysis (PCA) of soil macroinvertebrate data, combined with soil macromorphology features and soil chemical and physical properties: (**a**) Position of sampling sites on the plane defined by the first two PCA axes; ADE: Amazonian Dark Earth; REF: reference soils; OF: old forests; YF: young forests; AS: agricultural systems. Significance of Monte-Carlo test for soil type (ADE and REF) and land use systems (OF, YF and AS) *P* < 0.05. (**b**) Correlation circle representing the correlation between individual variables and the first two PCA axes. Blue arrows: macromorphological fractions (NAS=non-aggregated soil; PA=physical aggregates; RA=root aggregates; FA=fauna-produced aggregates, Pottery), total soil fauna density (number of ind. m^−2^), biomass (fresh biomass in g m^−2^) and overall morphospecies richness. (see Methods). Green arrows: density (no. ind. m^−2^) of ants, termites and earthworms. Red arrows: soil chemical properties (SB=sum of bases, CEC=cation exchange capacity, TC=total carbon, TN=total nitrogen) and particle size fractions (sand, silt, clay).

## Discussion

Our study found over 660 macroinvertebrate morphospecies in the 18 sites sampled in three Amazonian regions, including at least 70 new species of ecosystem engineers. We also found that although species richness is similar in ADE and REF soils, these two habitats harbour very different species pools, with few found in common to both habitats (Fig. 3b). Furthermore, although species rarefaction curves were still far from saturation using our current sampling effort, estimated richness showed similar trends, and showcased the wealth of species still to be discovered in both soils (Fig. 4). Finally, because these animals have been poorly represented in taxonomic surveys in Amazonia^22, 32–34^, and because ADEs had never been sampled before, we believe that these anthropogenic soils represent a major gap in the knowledge of Amazonian biodiversity. Although ADEs occupy only a small fraction of the Amazonian surface area^7^, they are scattered throughout the region^9, 35^, representing thousands of localized special habitats for species. The high β diversity values and species turnovers between different ADEs mean that each of these patches may be home to distinctive soil animal communities, including many new species, judging by the number of new ecosystem engineers found. Hence, ADEs represent an immense underground zoo, which could easily include thousands of species that have not yet been studied and/or classified.

Soil provides chemical and physical support for vegetation, and as millennia of human activities created ADEs in the Amazon, this generated patches of higher contents of nutrient and organic resources in a matrix of poorer soils^35^. The formation processes and human management of these soils results in distinct plant and microbial communities^6, 9, 13, 14^. Here we show that current soil animal abundance and diversity also reflect the impact of these ancient anthropogenic activities. The ADEs developed a different pool of species compared with REF soils. Similar biological selection processes probably occurred and are likely operating in other anthropogenic soils, either already created or being formed in various regions of the world (e.g., West Africa, Europe, Central America etc.)^36–38^. Studying the pathways to species selection (and possibly diversification) in ADEs and other anthropogenic soils requires further work, particularly expanding microbial and invertebrate biodiversity inventories. Fire may be one of the important factors to consider^39^: the anthropogenic alterations of ADE generally included frequent burning that led to the formation of highly stable charcoal^10^, and higher C and plant nutrient resources (Fig. 2)^12, 23^. Fire, in other contexts, has been documented to generate unique habitats that promote local biodiversity^40^.

The functional differences observed in biotic communities of ADEs also mean that these soils could provide different ecosystem services in the landscape. Higher earthworm populations and an improved soil structure mainly due to fauna-produced aggregates (as occurs in ADE) could positively affect primary productivity, litter decomposition and nutrient cycling^41^, pedogenetic processes^42^, and could help stabilize soil organic carbon in these soils^43^. These processes have been little studied, and merit further attention, both in forested and agriculturally managed ADE soils.

As archaeological sites, ADEs are protected by Brazilian law^44^, but throughout Amazonia they are intensively used for agricultural and horticultural purposes^35, 45, 46^. Soil macrofauna are threatened by modern land use change (particularly intensive annual cropping and livestock production), independently of the soil type. The biodiversity in both ADE and REF soils decreased with increasing environmental disturbance (Fig. 3a, Fig. 6), and negative impacts on populations of selected taxa were higher in ADE than in REF soils. Modern human activity has been associated with negative environmental impacts in the Amazon^3, 18^, but on the other hand, historical human footprints associated with ADEs appear to have “positive” effects on the Amazonian ecosystem^47^. For instance, we found that old forests on ADEs were the most biodiverse LUS.

Soil invertebrates are known to display high endemism^48^, and hence high β-diversity values, mainly due to their low dispersal ability^49^. Still, the high turnover rates between communities of ADE and REF soils suggest that ADEs may represent refuges for large numbers of specialist species that have been overlooked in previous work in the region^17, 18, 22, 32^, where ADEs were not targeted. This persistent anthropogenic footprint promotes biodiversity^50^ and modifies its distribution patterns in the Amazonian basin, making humans an endogenous part of the environment. This footprint is a prevailing driver in our study and as such, should be integrated into future ecological research in Amazonia. Finally, considering their distinctive below-ground communities, and the negative effect of modern land-use intensification, ADEs deserve special attention and management, in order to protect their biological resources and promote more sustainable uses of Amazonian soils^51^.

## Methods

### Study sites

The municipalities of Iranduba (IR) in Central Amazon, Belterra (BT) in Lower Amazon and Porto Velho (PV) in Southwestern Amazon, were chosen for this study (Fig. 1a). All sites have a tropical monsoon climate (Köppen’s Am), with a mean annual temperature of 24 °C and precipitation between 2,000 and 2,280 mm year^−1^ ^52^. In each region, paired sites with ADEs and nearby reference (REF) non-anthropogenic soils (Fig. 1b) were selected under different LUS (Fig. 1c): native secondary vegetation (*dense ombrophilous forest*) classified as old forest (OF) when >20 years old, or young forest (YF) when <20 years old, and agricultural systems (AS) of maize in IR, soybean in BT, and introduced pasture in PV. The REF sites were within a minimum distance of 150 m (soybean at BT) to a maximum distance of 1.3 km (pasture at PV) from the ADE sites, and maximum distance between paired sites within a region was 14 km (Embrapa sites to Tapajós National Forest sites in BT).

One of the OF in BT was at the Embrapa Amazônia Oriental Belterra Experiment Station, while the other one was at the Tapajós National Forest, a site of previous work on ADEs^39^. Both OFs at IR were at the Embrapa Amazônia Ocidental Caldeirão Experiment Station, and have been extensively studied in the past for soil fertility and pedogenesis^42, 53^, as well as microbial diversity^54, 55^. ADE formation in IR was estimated to have begun ∼1,050 - 950 years BP^56^ and at BT ∼530-450 years BP^57^. At PV, ADE formation began much earlier (∼6500 years BP)^8^.

The AS fields with annual crops were under continuous (at least 6 years) annual row cropping of maize (IR) and soybean (BT) and had been planted < 60 d prior to sampling, using conventional tillage (IR), or reduced tillage (BT). The crops received the recommended doses of inorganic fertilizers and pest management practices for each crop, which was planted using certified commercial seeds. The pastures at PV were around 9 (REF) and 12 yr old (ADE) and planted with *Brachiaria* (REF) and *Paspalum* (ADE) grasses. Soils at most REF sites were classified according to FAO^58^ as dystrophic Ferralsols and Acrisols (Supplementary Table 8), the two most common soil types in Amazonia^59^. At one YF site in PV, both ADE and REF soils were overlying a plinthic horizon and the REF soil was classified as a Plinthosol. All ADEs were classified as Pretic Clayic Anthrosols. with dark organic matter-rich surface soil horizons, generally >20 cm deep. All soils had greater than 50% clay and had either clay or heavy clayey texture. General details on the sampling sites chosen are provided in Supplementary Table 8.

### Soil macroinvertebrate sampling

We performed field sampling in April (IR) and May (BT) of 2015, and in late February/early March of 2016 (PV), at the end of the main rainy season, which is the best time to collect soil macroinvertebrates^60^. Soil and litter macrofauna were collected using the standard method recommended by the Tropical Soil Biology and Fertility (TSBF) Program of the United Nations Educational, Scientific and Cultural Organization (UNESCO)^61^, also considered by the International Organization for Standardization (ISO) as the appropriate method for evaluating soil macrofauna populations in the tropics^29^. At each sampling site, five sampling points were located within a 1 ha plot, at the corners and the centre of a 60 × 60m square, resulting in an “X” shaped sampling design (Supplementary Fig. 7). At each of these points, a soil monolith (25 × 25 cm up to 30 cm depth) was initially delimited with a 10 cm deep steel template, and then divided into surface litter and three 10 cm-thick layers (0-10, 10-20, 20-30 cm). Macroinvertebrates (i.e., invertebrates with > 2mm body width) were collected in the field by hand-sorting both the soil and litter, and were immediately fixed in 92% ethanol. Collected invertebrates were identified to species or genus level (earthworms, ants, termites), or sorted into morphospecies with higher taxonomic level assignations (e.g., order and/or family) for other groups.

### Additional samples for ecosystem engineers

We performed additional sampling for ecosystem engineers (earthworms, termites and ants), in order to better estimate their species richness, especially in forest sites where higher diversity is normally expected. Earthworms were collected at four additional cardinal points of the grid (Supplementary Fig. 7), hand-sorted from holes of similar dimensions as the TSBF monoliths, and preserved in 96% ethanol. Termites were sampled in five 10 m^2^ (2 × 5 m) plots (Supplementary Fig. 7) by manually digging the soil and looking for termitaria in the soil, as well as in the litter and on trees using a modification of the transect method^62^. The termite samples were taken in all OF and YF (except one of the REF YF at PV), but not in the agricultural fields (maize, soybean and pasture), as these tend to have very few termite colonies. Ants were sampled in 10 pitfall traps (300 ml plastic cups) set up as two 5-trap transects on the sides of each 1 ha plot (Supplementary Fig. 7), as well as in two traps to the side of each TSBF monolith (distant ∼5 m). Each cup was filled to a third of its volume with water, salt and detergent solution. The pitfall traps remained in the field for 48h. Pitfall traps were set up in only in the forest systems of IR and BT (not at PV). Termites and ants were preserved in 80% ethanol and the alcohol changed after cleaning the samples within 24 h. All the animals (earthworms, ants, termites) were identified to species level or morphospecies level (with genus assignations) by co-authors SWJ/MLCB (earthworms), AA (termites) and ACF/RMF (ants).

### Soil physical and chemical attributes

After hand-sorting the soil from each TSBF monolith, 2 to 3 kg samples were collected from each depth (0-10, 10-20, 20-30 cm) for chemical and soil particle size analysis, and while analysed separately, mean values were calculated over 0-30 cm depth. The following soil properties were assessed following standard methodologies: pH (CaCl_2_); Ca^2+^, Mg^2+^, Al^3+^ (KCl 1 mol L^−1^); K^+^ and P (Mehlich-1); total nitrogen (TN) and carbon (TC) using an element analyser (CNHS)^28^. Soil texture was obtained using the FAO soil texture triangle^58^, and base saturation and cation exchange capacity (CEC) were calculated using standard formulae^28^.

In order to assess functional differences induced by soil fauna activity in the ADE and REF soils, soil macromorphology samples were taken 2 m from each monolith (Supplementary Fig. 7) using a 10 × 10 × 10 cm metal frame. The collected material was separated into different fractions including: living invertebrates, litter, roots, pebbles, pottery shards, charcoal (biochar), non-aggregated/loose soil, physical aggregates, root-associated aggregates, and fauna-produced aggregates using the method of Velásquez et al.^30^. Each fraction was oven dried at 60°C for 24h and weighed. This method allows estimating the relative contribution of soil macrofauna, roots and soil physical processes to soil macroaggregation^30^ and structure, which determines the delivery of several important soil-based ecosystem services^63^.

### Treatment of soil fauna data

Density (number of individuals) and biomass of the soil macrofauna surveyed using the TSBF method were extrapolated per square meter considering all depths evaluated. Density and biomass of immature forms of insects (nymphs and larvae) were grouped in the respective taxonomic group. The following taxonomic groups, representing 2% or less of total density were grouped as “Others”: Araneae, Hemiptera, Orthoptera, Diptera (larvae), Gastropoda, Dermaptera, Isopoda, Blattaria, Scorpionida, Opiliones, Lepidoptera (larvae), Uropygi, Solifuga, Thysanoptera, Geoplanidae, Neuroptera (larvae), Hirudinea and Embioptera. To calculate the beta (β) diversity index we removed singleton species (species represented by single individuals, i.e., one individual among all the 9,380 individuals collected).

### Statistical analyses

To compare species diversity between ADE and REF, we plotted rarefaction and extrapolation curves using the iNEXT^64^ package for total macroinvertebrate, ant, termite and earthworm species diversity, using the number of TSBF monolith samples as a measure of sampling effort intensity. The same procedure was used for all earthworm data (9 samples per site), termite data obtained from both the 10-m^2^ plots and TSBF monoliths, and ant data obtained from both pitfall traps and TSBF monoliths.

We used the betapart package^65^ in R to decompose β-diversity (calculated using the Sørensen dissimilarity index) into its Turnover (Simpson index of dissimilarity) and Nestedness components using all soil+litter macroinvertebrate, ant, termite and earthworm data from monolith samples. The average β-diversity was calculated to highlight LUS effect, by comparing all LUS (OF, YF and AS) within each soil type (REF and ADE) and region. The soil type effect was assessed comparing the diversity between REF and ADE soils within each LUS in each region. To identify the effect of geographical distance on species turnover we also calculated the average β-diversity among the three replicates of each LUS within each soil type.

Due to non-normal distribution of both the faunal variables (i.e., density and biomass of invertebrates collected using the TSBF method) and soil properties, we used General Linear Models (GLM) to adjust data to other probability distributions. The best adjustment was quasi-Poisson (overdispersion) and Gamma for invertebrate density and biomass, respectively. Soil chemical properties were adjusted in Gamma distribution but particle size fractions could not be adjusted. ANOVA tests were performed with the multcomp package^66^ of R, adopting a factorial design with the following factors: soil type (ADE and REF) and LUS (old forests, young forests and agricultural systems). When factor interactions were significant (*P* < 0.05), the data were analysed comparing the effects of soil type within the LUS and the effects of LUS within each soil type. Significant differences were tested using Tukey’s test at 95% probability (*P* < 0.05) for GLM, or with non-parametric Kruskal-Wallis tests when data could not be adjusted with GLM.

A Principal Component Analysis (PCA) was performed using the density of earthworms, termites, ants and overall (total) soil fauna density and biomass, together with the results of five variables from soil micromorphology (non-aggregated soil, pottery shards and fauna, root and physical aggregates) and ten variables from soil chemical and textural analyses (pH, Al^3+^, P, SB, T, TC, TN, and sand, silt and clay fractions). The significance of the PCA model (soil type and LUS) was assessed using Monte Carlo test permutations (*P* < 0.05), using the ADE-4 package^67^ for R.

## Reporting Summary

Further information on experimental design is available in the Nature Research Reporting Summary linked to this article.

## Data availability

All the data generated and analysed during this study are included in this published article and its supplementary information files. Raw data are available from the authors upon reasonable request.

## Acknowledgements

The study was supported by the Newton Fund and Fundação Araucária (grant Nos. 45166.460.32093.02022015, NE/N000323/1), Natural Environment Research Council (NERC) UK (grant No. NE/M017656/1), a European Union Horizon 2020 Marie-Curie fellowship to LC (MSCA-IF-2014-GF-660378) and DWGS (No. 796877), by CAPES scholarships to WCD, ACC, TF, RFS, AF, LM, HSN, TS, AM and RSM (PVE A115/2013), Araucaria Foundation scholarships to LM, AS, ACC and ES, Post-doctoral fellowships to DWGS (NERC grant NE/M017656/1) and ES (CNPq No. 150748/2014-0), PEER (Partnerships for Enhanced Engagement in Research Science Program) NAS/USAID award number AID-OAA-A-11-0001 - project 3-188 to RMF and by CNPq grants, scholarships and fellowships to ACF, GGB, RF, SWJ, CRC, EGN and PL (Nos. 140260/2016-1, 307486/2013- 3, 302462/2016-3, 401824/2013-6, 303851/2015-5, 307179/2013-3, 400533/2014-6). We thank INPA, UFOPA, Embrapa Rondônia, Embrapa Amazônia Ocidental and Embrapa Amazônia Oriental and their staff for logistical support and the farmers for access to and permission to sample on their properties. Sampling permit for Tapajós National Forest was granted by ICMBio.

## Author Contributions

LC, PK and GGB conceived the study and planned the field work. WCD, ACC, AF, AA, MLCB, SWJ, ES, GCM, RSM, DWGS, PL, EV, AZ, RB, STC, AWM, AS, TF, RFS, TD, HSN, CPV, AP, AM, PT, TS, PK, GGB and LC carried out the field work. WCD, ACC, ES, LC and GGB analysed all the data. LC, PK and GGB conceived the study and planned the field work. WCD, GGB and LC wrote the manuscript, and all authors contributed to and approved the final version. LC and GGB have jointly co-supervised the work and both share the senior authorship of the work.

## Additional information

Supplementary Information accompanies this paper

