## Supplemental File for "A “Dirty” Footprint: Anthropogenic Soils Promote Biodiversity in Amazonian Rainforests"

### Supplementary Information

**Contents:**

Supplementary tables

**Supplementary Table 1**. Soil analyses from the topsoil layers

**Supplementary Table 2**. Morphospecies richness of the main soil macroinvertebrate taxa and groups

**Supplementary Table 3**. Number of singleton, doubleton, rare, and abundant species/morphospecies

**Supplementary Table 4**. Soil macroinvertebrate density and biomass

**Supplementary Table 5**. Aggregate fractions from the micromorphology samples

**Supplementary Table 6**. Effects of land-use systems (LUS) and soil type (REF and ADE) on β-diversity

**Supplementary Table 7**. Effects of region on Beta diversity

Supplementary Figures

**Supplementary Figure 1**. Venn charts of total macroinvertebrate species richness

**Supplementary Figure 2**. Venn charts of ant species richness

**Supplementary Figure 3**. Venn charts of termite species richness

**Supplementary Figure 4**. Venn charts of earthworm species richness

**Supplementary Figure 5**. Morphospecies rarefaction and extrapolation curves

**Supplementary Figure 6**. Sampling effort coverage

**Supplementary Figure 7.** Scheme used for soil and fauna sampling

**Supplementary Table 1. Soil analyses from the topsoil layers** (0-30 cm depth) in reference (REF) and Amazonian Dark Earth (ADE) soils under each of the land-use systems (OF: old forests, YF: young forests, AS: agricultural systems). Values represent means from the three study regions.

| Soil type | Land use | Al[1] | K[1] | CEC[1] | Base saturation[1] | Total N[1] | Sand[2] | Silt[2] | Clay[2] | Texture class (FAO) |
|---|---|---|---|---|---|---|---|---|---|---|
| | | - - - - - - - - - - - - - - - - $cmol_c\ kg^{-1}$ - - - - - - - - - - - - - - - - | | | | - - - - - - - - - - - - - - - - - % - - - - - - - - - - - - - - - - - | | | | |
| REF | OF | 2.3±0.14Aa | 0.06±0.01Ab | 3.16±0.22Ba | 7.6±2.1Ba | 0.22±0.01Aab | $23±7^{ns}$ | 15±2Ab | 62±5Aa | Heavy clay |
| | YF | 2.8±0.09Aa | 0.05±0.0Aab | 2.89±0.09Ba | 1.2±0.1Bb | 0.27±0.03Aa | 26±4 | 22±2Aa | 52±2Ab | Clay |
| | AS | 1.6±0.22Aa | 0.09±0.01Aa | 3.16±0.34Ba | 19.5±5.7Ba | 0.21±0.01Ab | 21±4 | 19±9Aa | 60±3Aa | Heavy clay |
| ADE | OF | 0.5±0.20Ba | 0.07±0.01Ab | 8.76±0.78Aa | 55.1±4.9Aa | 0.28±0.02Aab | 23±5 | 17±2Ab | 60±4Aa | Heavy clay |
| | YF | 0.6±0.13Ba | 0.10±0.01Aab | 8.85±1.0Aa | 49.2±4.6Aa | 0.36±0.03Aa | 27±2 | 23±2Aa | 50±2Ab | Clay |
| | AS | 0.4±0.11Ba | 0.12±0.02Aa | 6.95±0.69Aa | 50.3±4.8Aa | 0.25±0.01Ab | 19±3 | 22±1Aa | 59±2Aa | Clay |

*Upper case letters compare soil categories (ADE vs. REF), within each land-use system, while lower-case letters compare land use systems within the same soil type (ADE or REF). Different letters mean significant differences resulting from GLM or Kruskal-Wallis non-parametric tests.
[1]GLM; [2]KW.

**Supplementary Table 2. Morphospecies richness of the main soil macroinvertebrate taxa and groups.** Number of species/morphospecies of each of the major soil macroinvertebrate taxa collected in the soil monoliths (litter and 0-30 cm), in each of the three main land-use systems (OF: old forests, YF: young forests, AS: agricultural systems) in both reference (REF) and Amazonian Dark Earth (ADE) soils (sum of all three regions), and total observed richness of each taxon collected over all samples. Total numbers do not always represent sum of all species in each land-use system due to shared species/morphospecies.

| Group | Number of morphospecies collected (soil monoliths) | | | | | | | | Overall observed richness |
|---|---|---|---|---|---|---|---|---|---|
| | REF | | | | ADE | | | | |
| | OF | YF | AS | Total | OF | YF | AS | Total | |
| **Earthworms** | 12 | 9 | 4 | 22 | 11 | 10 | 7 | 22 | 32 |
| **Termites** | 13 | 20 | 9 | 32 | 9 | 12 | 0 | 16 | 37 |
| **Ants** | 58 | 41 | 18 | 94 | 58 | 35 | 24 | 93 | 154 |
| **Beetles** | 16 | 21 | 15 | 47 | 25 | 11 | 12 | 43 | 78 |
| **Spiders** | 22 | 15 | 10 | 47 | 21 | 15 | 10 | 44 | 86 |
| **Millipedes** | 12 | 9 | 7 | 25 | 14 | 23 | 5 | 37 | 53 |
| **Centipedes** | 8 | 8 | 2 | 14 | 10 | 5 | 3 | 11 | 17 |
| **True bugs** | 9 | 5 | 11 | 24 | 8 | 11 | 4 | 21 | 42 |
| **Cockroaches** | 11 | 7 | 2 | 20 | 8 | 12 | 1 | 20 | 34 |
| **Opiliones** | 6 | 8 | 0 | 14 | 5 | 5 | 0 | 9 | 21 |
| **Isopods** | 6 | 6 | 1 | 11 | 7 | 5 | 2 | 13 | 21 |
| **Snails** | 3 | 2 | 2 | 7 | 7 | 3 | 2 | 12 | 17 |
| **Others*** | 20 | 14 | 10 | 42 | 25 | 14 | 7 | 44 | 75 |

*This group includes Orthoptera, Diptera (larvae), Dermaptera, Scorpionida, Lepidoptera (larvae), Uropygi, Solifuga, Thysanoptera, Geoplanidae, Neuroptera (larvae), Hirudinea and Embioptera

**Supplementary Table 3. Number of singleton, doubleton, rare, and abundant species/morphospecies** of all soil macroinvertebrate taxa and of selected macrofauna taxa collected in reference (REF) and Amazonian Dark Earth (ADE) soils (sum of all three regions and land use systems). Rare species represent taxa with fewer than 10 ind. over all samples. Non-rare and abundant species represent taxa with ≥10 ind. over all samples. Unique species were found in either ADE or REF; shared species were found in both soil categories. Species more abundant were considered to be those with quantities at least three times greater in ADE or REF soils, respectively. Locally distributed species were those found only in one region but in more than one land-use system. Widely distributed species were those found in more than one region.

| Soil type | Taxon | Singletons | Doubletons | | Rare | | Non-rare or Abundant | | | | | | |
|---|---|---|---|---|---|---|---|---|---|---|---|---|---|
| | | | Unique | Shared | Unique | Shared | Unique | Shared | More abundant in REF | Locally distributed (Total) | More abundant in REF | Widely distributed (Total) | More abundant in REF |
| REF | | | | | | | | | | | | | |
| | Ants | 29 | 8 | 2 | 15 | 11 | 8 | 20 | 6 | 12 | 2 | 16 | 4 |
| | Termites | 6 | 3 | - | 7 | 3 | 8 | 5 | 2 | 9 | 1 | 4 | 3 |
| | Earthworms | 5 | - | - | 5 | 2 | - | 10 | - | 7 | - | 3 | - |
| | Beetles | 28 | 4 | 3 | 4 | 4 | - | 4 | 1 | 1 | 1 | 3 | - |
| | Millipedes | 5 | 3 | - | 8 | 4 | - | 5 | 2 | 4 | 2 | 1 | - |
| | Total invertebrates | 171 | 34 | 14 | 61 | 42 | 19 | 56 | 13 | 48 | 4 | 27 | 9 |
| ADE | | | | | | | | | More abundant in ADE | | More abundant in ADE | | More abundant in ADE |
| | Ants | 26 | 10 | 2 | 17 | 11 | 8 | 20 | 7 | 9 | 2 | 19 | 5 |
| | Termites | - | 2 | - | 2 | 3 | 1 | 5 | 1 | 2 | - | 4 | 1 |
| | Earthworms | 2 | 2 | - | 3 | 2 | 2 | 10 | 7 | 1 | 5 | 3 | 2 |
| | Beetles | 20 | 4 | 3 | 4 | 4 | 1 | 4 | 1 | 1 | - | 3 | 1 |
| | Millipedes | 14 | 6 | - | 6 | 4 | 2 | 5 | 1 | 6 | 1 | 1 | - |
| | Total invertebrates | 157 | 43 | 14 | 52 | 42 | 18 | 56 | 19 | 44 | 9 | 30 | 10 |

**Supplementary Table 4. Soil macroinvertebrate density and biomass.** Mean density (Den.; number of individuals $m^{-2}$) and biomass (Bio.; fresh mass in g $m^{-2}$) of the main soil macroinvertebrate taxa collected in each of the land-use systems (OF: old forests, YF: young forests, AS: agricultural systems) studied in reference (REF) and Amazonian Dark Earth (ADE) soils. Different lower-case letters indicate significant differences between land-use systems within the same soil type, while different upper-case letters indicate significant differences between soil categories within each land-use system. Significance determined using GLMs or Kruskal-Wallis (KW) non-parametric tests.

| GROUPS | | REF | | | ADE | | |
|---|---|---|---|---|---|---|---|
| | | OF | YF | AS | OF | YF | AS |
| **Earthworms[1]** | Den. | 152.5±28.2Aa | 60.8±17.5Ba | 44.8±12.4Bb | 193.1±29.7Aa | 268.8±35.2Aa | 234.7±58.5Aa |
| | Bio. | 19.34±4.48Aa | 10.30±3.46Bab | 3.58±1.09Bb | 13.39±3.15Aa | 27.34±5.11Aa | 18.07±7.99Aa |
| **Termites[2]** | Den. | 1758.9±989.2Aa | 1057.1±365.2Aa | 1241.6±858Aa | 141.9±123.9Bab | 168.5±80.3Ba | 1.1±1.0Bb |
| | Bio. | 3.87±2.58Aa | 2.58±0.99Aa | 2.11±0.96Aa | 0.17±0.14Bab | 0.34±0.18Ba | 0.00±0.0Bb |
| **Ants[1]** | Den. | 476.8±158.1Aa | 210.1±69.4Aa | 137.6±60.3Aa | 416.0±111.9Aa | 231.5±80.3Aa | 533.3±316.3Aa |
| | Bio. | 0.68±0.21Aa | 0.41±0.11Aa | 0.14±0.04Ab | 0.76±0.26Aa | 0.54±0.22Aa | 0.47±0.19Aa |
| **Ecosystem engineers[1]** | Den. | 2388.3±956.4Aa | 1328±362.2Aa | 1424±367.7Aa | 750.9±193.9Ba | 668.9±172.7Ba | 769.1±198.6Ba |
| | Bio. | 23.89±5.66Aa | 13.28±3.41Bab | 5.83±1.41Ab | 14.32±3.08Aa | 28.22±5.03Aa | 18.54±7.96Aa |
| **Beetles[1]** | Den. | 52.3±9.6Ba | 58.7±11.2Aa | 57.6±16.4Aa | 137.6±33.0Aa | 52.3±14.0Ab | 21.3±6.0Bb |
| | Bio. | 1.73±1.07Ba | 2.60±1.18Aa | 0.58±0.18Aa | 3.11±0.96Aa | 0.58±0.20Ab | 0.21±0.11Ab |
| **Millipedes[2]** | Den. | 25.6±8.8Aa | 24.5±6.6Ba | 52.3±20.7Aa | 37.3±6.0Aab | 97.1±32.1Aa | 13.9±5.2Ab |
| | Bio. | 1.02±0.50Aa | 0.24±0.07Ba | 0.75±0.29Aa | 0.63±0.21Aab | 3.50±1.54Aa | 0.24±0.10Ab |
| **Centipedes[2]** | Den. | 50.1±11.6Aa | 38.4±9.0Aa | 4.3±1.8Ab | 67.2±10.3Aa | 58.7±20.4Aab | 19.2±8.2Ab |
| | Bio. | 0.49±0.23Aa | 0.29±0.09Aa | 0.03±0.01Ab | 0.27±0.05Aa | 0.31±0.11Aab | 0.11±0.06Ab |
| **Others[2]*** | Den. | 103.5±13.5Ba | 58.7±10.3Aa | 106.7±30.4Aa | 168.5±28.7Aa | 113.1±28.7Aab | 68.3±22.1Ab |
| | Bio. | 6.47±2.29Aa | 0.34±0.10Bb | 0.94±0.37Ab | 3.82±0.78Aa | 1.23±0.34Ab | 0.61±0.20Ab |
| **Total[2]** | Den. | 2619.7±951.4Aa | 1508.3±358.5Aa | 1644.8±851Aa | 1161.6±187.5Aa | 989.9±156.4Aa | 891.7±318Aa |
| | Bio. | 33.59±6.97Aa | 16.75±3.69Bab | 8.13±1.35Ab | 22.15±3.65Aa | 33.84±5.44Aa | 19.72±7.82Aa |

[1]GLM; [2]KW; *This group includes Orthoptera, Diptera (larvae), Dermaptera, Hemiptera, Isopoda, Blattaria, Gastropoda, Lepidoptera (larvae), Uropygi, Solifuga, Opiliones, Scorpionida, Thysanoptera, Geoplanidae, Neuroptera (larvae), Hirudinea, and Embioptera

**Supplementary Table 5. Aggregate fractions from the micromorphology samples.** Mean relative biomass (%) of the different aggregate fractions found in the soil macromorphology samples (0-10 cm) in each of the land-use systems (OF: old forests, YF: young forests, AS: agricultural systems) in reference (REF) and Amazonian Dark Earth (ADE) soils. Fractions measured were biogenic aggregates produced by ecosystem engineers (Fauna), rhizosphere aggregates (Root), physical aggregates (Physical), non-macroaggregated loose soil particles and unidentified aggregates less than 5 mm in size (NAS), coarse organic material such as leaves, roots, seeds, and woody pieces (Organic materials), soil invertebrates (Invertebrates), pottery shards (Pottery), stones and charcoal. Different lower-case letters indicate significant differences between land-use systems within the same soil type, while upper-case letters compare soil categories within each land-use system. Mean comparisons performed using ANOVA, GLMs or Kruskal-Wallis (KW) non-parametric tests.

| Aggregate fraction | REF | | | ADE | | |
|---|---|---|---|---|---|---|
| | OF | YF | AS | OF | YF | AS |
| **Fauna[1]** | 43.6±5.3Ba | 22.6±1.8Bb | 14.2±1.1Bb | 48.7±5.4Aa | 28.8±0.4Ab | 23.6±0.2Ab |
| **Root[2]** | 4.4±1.1Aab | 8.3±1.4Aa | 5.9±2.1Ab | 5.1±1.5Aab | 10.5±2.4Aa | 8.9±3.4Ab |
| **Physical[2]** | 0.7±0.5Ac | 7.5±1.6Ab | 27.5±3.6Aa | 0.0±0.0Ac | 9.6±2.6Ab | 20.3±2.7Aa |
| **NAS*** | 41.7±5.4Ab | 52.5±2.1Aa | 50.4±3.0Aab | 36.5±2.8Bb | 45.9±2.9Ba | 42.7±4.2Bab |
| **Organic materials[1]** | 8.3±1.0Aa | 4.0±1.0Ab | 1.9±0.4Ab | 5.8±1.0Aa | 2.0±0.4Ab | 1.9±0.3Ab |
| **Invertebrates[2]** | 1.0±0.4$^{ns}$ | 0.0±0.0 | 0.1±0.4 | 1.4±0.8 | 0.0±0.0 | 0.2±0.1 |
| **Pottery[2]** | 0.2±0.2$^{ns}$ | 0.0±0.0 | 0.0±0.0 | 1.8±0.6 | 1.0±0.3 | 2.3±0.9 |
| **Stones[2]** | 0.0±0.0$^{ns}$ | 4.4±2.1 | 0.0±0.0 | 0.5±0.3 | 2.1±2.0 | 0.1±0.1 |
| **Charcoal[2]** | 0.20±0.2$^{ns}$ | 0.59±0.3 | 0.00±0.0 | 0.003±0.0 | 0.06±0.1 | 0.01±0.0 |

[1]GLM; [2]KW; *ANOVA

**Supplementary Table 6. Effects of land-use systems (LUS) and soil type (REF and ADE) on β-diversity** (without singletons), species turnover rates and nestedness of total soil macrofauna (339 morphospecies), ants, termites and earthworm communities. Richness values used for the calculations are from the soil monoliths (TSBF). REF: Reference soil, ADE: Amazonian Dark Earth, OF: old forests, YF: young forests, AS: agricultural systems.

| | Max div. ($\beta_{Sorensen}$) | Turnover ($\beta_{Simpson\ dis.}$) | Nestedness |
|---|---|---|---|
| **Macroinvertebrates** | | | |
| LUS effect | | | |
| on REFs | 0.86 | 0.79 | 0.07 |
| on ADEs | 0.83 | 0.74 | 0.09 |
| Soil effects | | | |
| in OF | 0.72 | 0.70 | 0.02 |
| in YF | 0.70 | 0.67 | 0.03 |
| in AS | 0.77 | 0.71 | 0.06 |
| **Ants** | | | |
| LUS effect | | | |
| on REFs | 0.86 | 0.72 | 0.14 |
| on ADEs | 0.83 | 0.75 | 0.08 |
| Soil effects | | | |
| in OF | 0.80 | 0.78 | 0.28 |
| in YF | 0.80 | 0.74 | 0.06 |
| in AS | 0.78 | 0.67 | 0.11 |
| **Termites** | | | |
| LUS effect | | | |
| on REFs | 0.81 | 0.65 | 0.16 |
| on ADEs | 0.81 | 0.39 | 0.42 |
| Soil effects | | | |
| in OF | 0.79 | 0.72 | 0.07 |
| in YF | 0.64 | 0.53 | 0.12 |
| in AS | - | - | - |
| **Earthworms** | | | |
| LUS effect | | | |
| on REFs | 0.90 | 0.84 | 0.06 |
| on ADEs | 0.72 | 0.62 | 0.10 |
| Soil effects | | | |
| in OF | 0.34 | 0.11 | 0.23 |
| in YF | 0.50 | 0.38 | 0.13 |
| in AS | 0.78 | 0.67 | 0.11 |

**Supplementary Table 7. Effects of region on Beta diversity** (without singletons), species turnover rates and nestedness of total soil macrofauna (339 morphospecies), ants, termites and earthworm communities, among each land-use system (OF: old forest; YF: young forest; AS: agricultural systems) within each soil type (REF and ADE). Richness values used for the calculations are from the soil monoliths (TSBF). REF: Reference soil, ADE: Amazonian Dark Earth.

| | Max div. ($\beta_{Sorensen}$) | Turnover ($\beta_{Simpson\ dis.}$) | Nestedness |
|---|---|---|---|
| **Macroinvertebrates** | | | |
| Region effect | | | |
| OF - REF | 0.85 | 0.81 | 0.04 |
| ADE | 0.82 | 0.81 | 0.01 |
| YF - REF | 0.86 | 0.81 | 0.05 |
| ADE | 0.85 | 0.79 | 0.06 |
| AS - REF | 0.91 | 0.90 | 0.01 |
| ADE | 0.90 | 0.86 | 0.04 |
| **Ants** | | | |
| Region effect | | | |
| OF - REF | 0.81 | 0.71 | 0.10 |
| ADE | 0.83 | 0.80 | 0.03 |
| YF - REF | 0.89 | 0.82 | 0.07 |
| ADE | 0.80 | 0.73 | 0.07 |
| AS - REF | 0.81 | 0.66 | 0.15 |
| ADE | 0.75 | 0.68 | 0.07 |
| **Termites** | | | |
| Region effect | | | |
| OF - REF | 0.82 | 0.72 | 0.10 |
| ADE | 0.83 | 0.75 | 0.08 |
| YF - REF | 0.71 | 0.63 | 0.08 |
| ADE | 0.81 | 0.77 | 0.04 |
| AS - REF | 0.88 | 0 | 0.88 |
| ADE | - | - | - |
| **Earthworms** | | | |
| Region effect | | | |
| OF - REF | 0.69 | 0.60 | 0.09 |
| ADE | 0.75 | 0.70 | 0.05 |
| YF - REF | 1 | 1 | 0 |
| ADE | 0.77 | 0.75 | 0.01 |
| AS - REF | 1 | 1 | 0 |
| ADE | 1 | 1 | 0 |

**Supplementary Table 8. General geographic, soil and land use information on the sampling sites.** Land use system, age of modern human intervention, soil type, soil category according to WRV/FAO (2015) and location of the sites studied in the three regions of Brazilian Amazonia. Dark green colour represents old forests, pale green colour represents young forests and yellow colour represents agricultural systems.

| Region | State | Land use | Human intervention[1] | Soil type[2] | Soil category (WRB) | Coordinates |
|---|---|---|---|---|---|---|
| Iranduba | AM | Old Forest | > 20 years old | REF | Xanthic Dystric Acrisol | 3°14'49.00"S, 60°13'30.71"W |
| | | Old Forest | > 20 years old | ADE | Pretic Clayic Anthrosol | 3°15'11.05"S, 60°13'45.03"W |
| | | Young Forest | < 20 years old | REF | Xanthic Dystric Acrisol | 3°13'34.47"S, 60°16'23.60"W |
| | | Young Forest | < 20 years old | ADE | Pretic Clayic Anthrosol | 3°13'49.23"S, 60°16'7.43"W |
| | | Agricultural | Current | REF | Xanthic Dystric Acrisol | 3°13'31.31"S, 60°16'29.18"W |
| | | Agricultural | Current | ADE | Pretic Clayic Anthrosol | 3°13'46.13"S, 60°16'7.32"W |
| Belterra | PA | Old Forest | > 20 years old | REF | Xanthic Dystric Ferralsol | 2°47'4.59"S, 54°59'53.28"W |
| | | Old Forest | > 20 years old | ADE | Pretic Clayic Anthrosol | 2°47'3.25"S, 54°59'59.77"W |
| | | Old Forest | > 20 years old | REF | Xanthic Dystric Acrisol | 2°41'13.90"S, 54°55'3.30"W |
| | | Old Forest | > 20 years old | ADE | Pretic Clayic Anthrosol | 2°41'7.18"S, 54°55'7.11"W |
| | | Agricultural | Current | REF | Xanthic Dystric Acrisol | 2°41'3.56"S, 54°55'12.75"W |
| | | Agricultural | Current | ADE | Pretic Clayic Anthrosol | 2°41'3.79"S, 54°55'7.90"W |
| Porto | RO | Young Forest | < 20 years old | REF | Xanthic Dystric Plinthosol | 8°52'11.50"S, 64° 3'18.16"W |
| Velho | | Young Forest | < 20 years old | ADE | Pretic Clayic Anthrosol | 8°51'51.92"S, 64°03'48.03"W |
| | | Young Forest | < 20 years old | REF | Xanthic Dystric Ferralsol | 8°50'49.52"S, 64° 3'59.20"W |
| | | Young Forest | < 20 years old | ADE | Pretic Clayic Anthrosol | 8°51'1.18"S, 64° 4'3.07"W |
| | | Agricultural | Current | REF | Xanthic Dystric Ferralsol | 8°52'35.30"S, 64°03'58.58"W |
| | | Agricultural | Current | ADE | Pretic Clayic Anthrosol | 8°51'56.53"S, 64°03'40.67"W |

[1]Age of modern human disturbance (land management);
[2]REF –reference soil, ADE – Amazonian Dark Earth

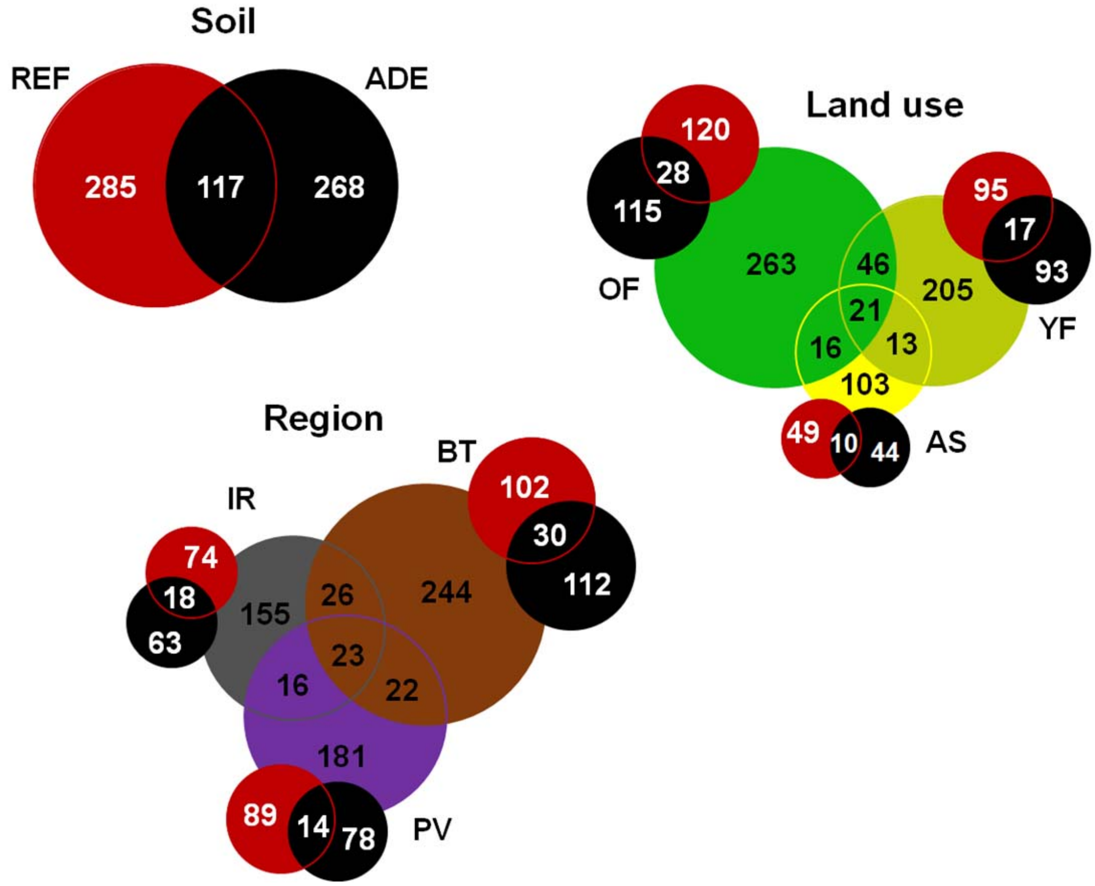

**Supplementary Figure 1. Venn charts of total macroinvertebrate species richness** (from soil monolith samples), and overlaps according to soil (REF: Reference soil, ADE: Amazonian Dark Earth), to geographic region (IR: Iranduba; BT: Belterra; PV: Porto Velho), and land-use systems (OF: old forest; YF: young forest; AS: agricultural systems) (ADE in black, REF in red). Numbers for species richness in REF (in red) and ADE (in black) soils are only of the unique species in each region and land-use system.

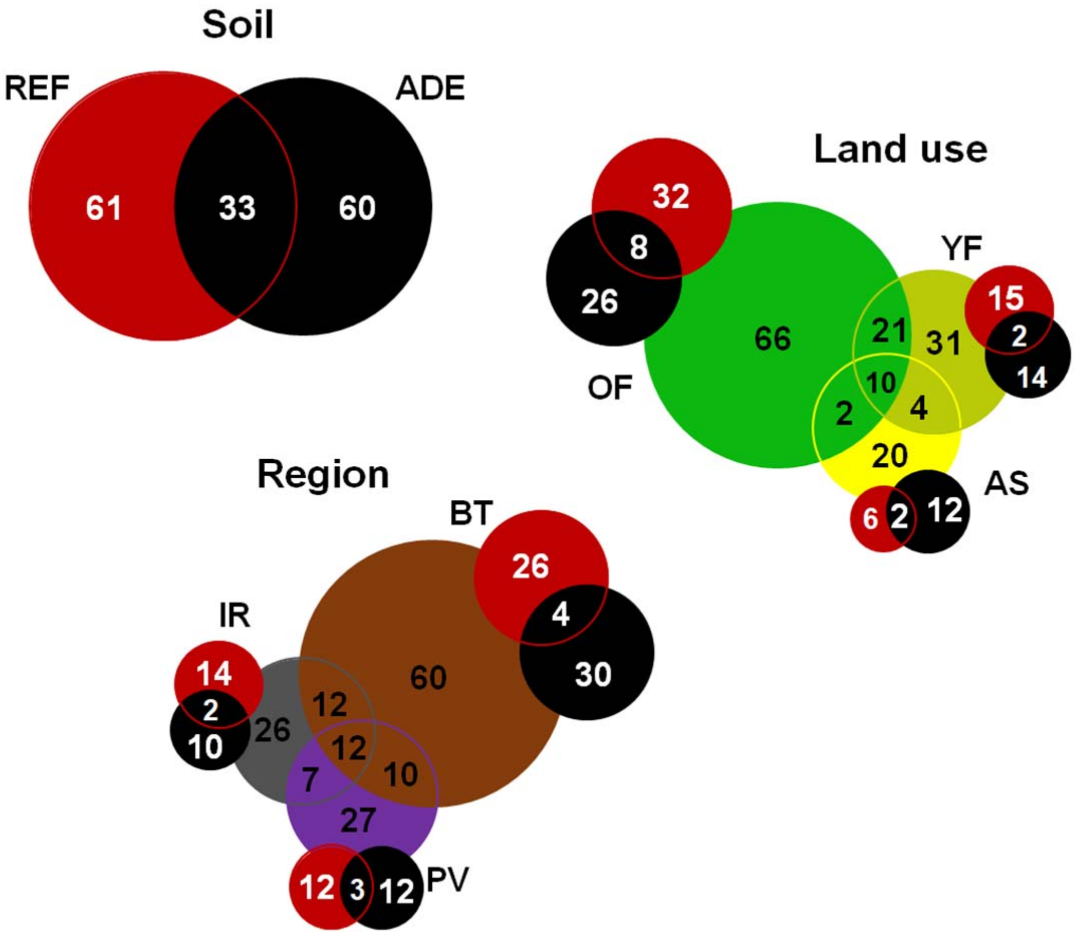

**Supplementary Figure 2. Venn charts of ant species richness** (from soil monolith samples), and overlaps according to soil (REF: Reference soil, ADE: Amazonian Dark Earth), to geographic region (IR: Iranduba; BT: Belterra; PV: Porto Velho), and land-use systems (OF: old forest; YF: young forest; AS: agricultural systems) (ADE in black, REF in red). Numbers for species richness in REF (in red) and ADE (in black) soils are only of the unique species in each region and land-use system.

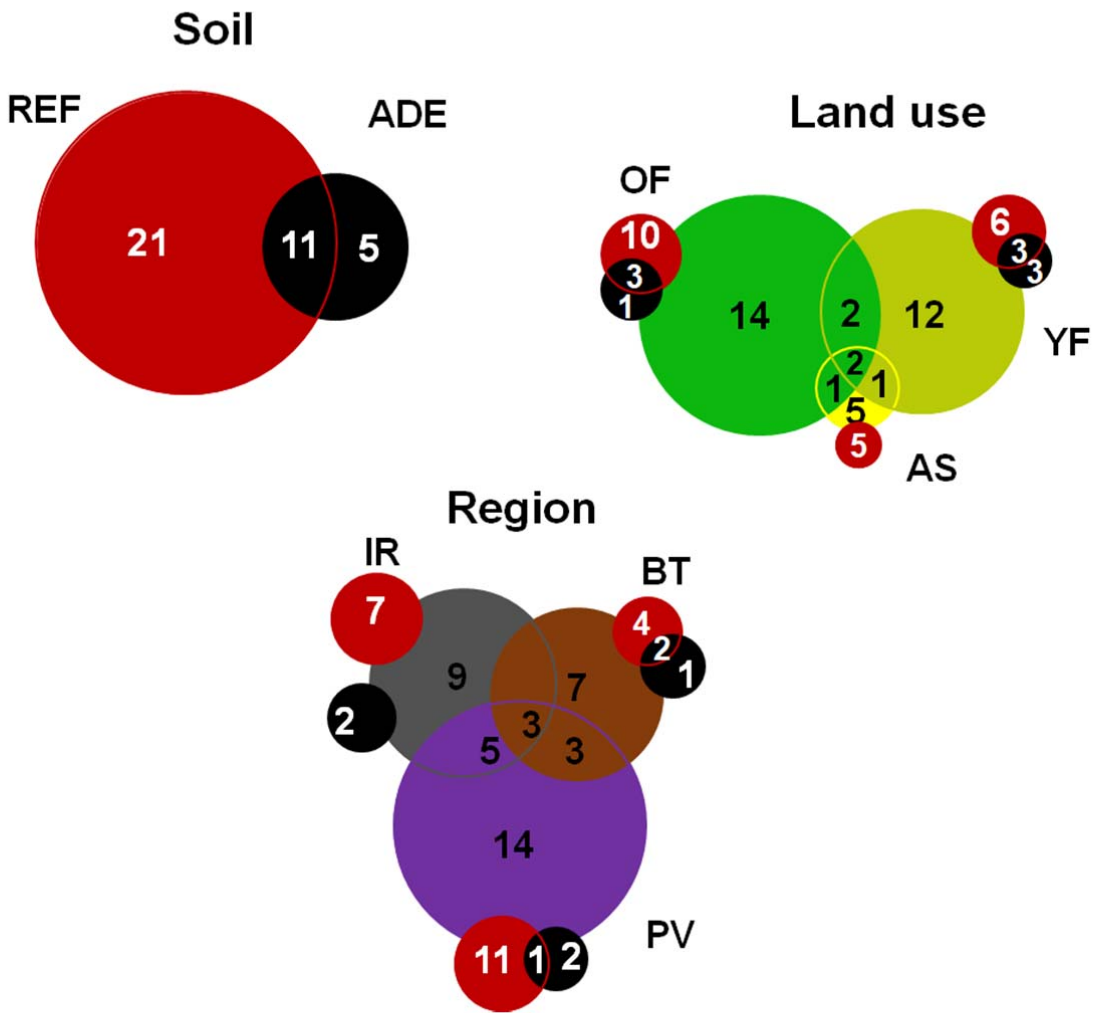

**Supplementary Figure 3. Venn charts of termite species richness** (from soil monolith samples), and overlaps according to soil (REF: Reference soil, ADE: Amazonian Dark Earth), to geographic region (IR: Iranduba; BT: Belterra; PV: Porto Velho), and land-use systems (OF: old forest; YF: young forest; AS: agricultural systems) (ADE in black, REF in red). Numbers for species richness in REF (in red) and ADE (in black) soils are only of the unique species in each region and land-use system.

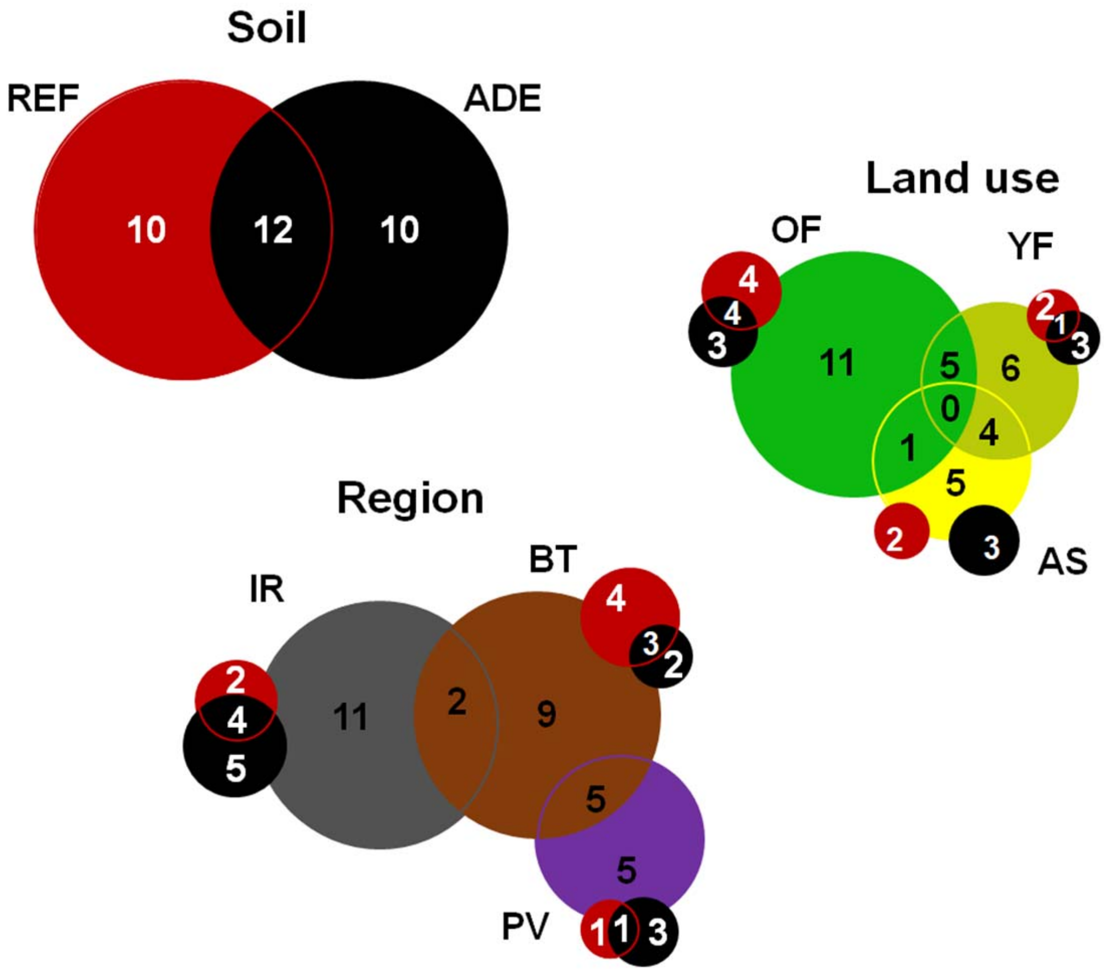

**Supplementary Figure 4. Venn charts of earthworm species richness** (from soil monolith samples), and overlaps according to soil (REF: Reference soil, ADE: Amazonian Dark Earth), to geographic region (IR: Iranduba; BT: Belterra; PV: Porto Velho), and land-use systems (OF: old forest; YF: young forest; AS: agricultural systems) (ADE in black, REF in red). Numbers for species richness in REF (in red) and ADE (in black) soils are only of the unique species in each region and land-use system.

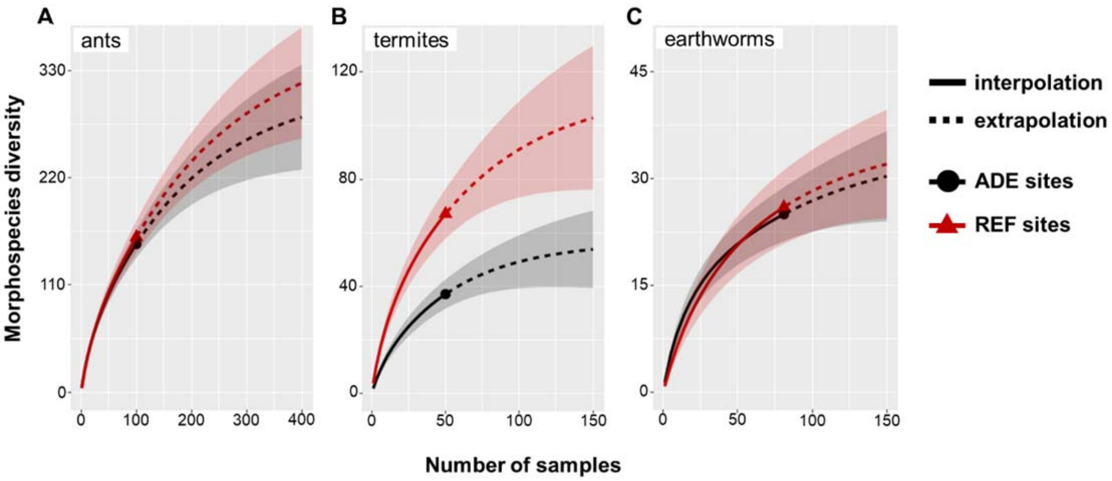

**Supplementary Figure 5. Morphospecies rarefaction and extrapolation curves**, showing how morphospecies numbers increase in both ADE and REF soils depending on sampling intensity (number of samples) for: (**a**) ants collected in pitfall traps + TSBF in Iranduba and Belterra under old and young forests, (**b**) termites in TSBF samples + 10 $m^2$ plots in old and young forests (except YF1 in PV), and (**c**) earthworms from all (n=9 per plot) samples over all sites. Dark grey and red areas represent 95% confidence intervals. REF: Reference soil, ADE: Amazonian Dark Earth

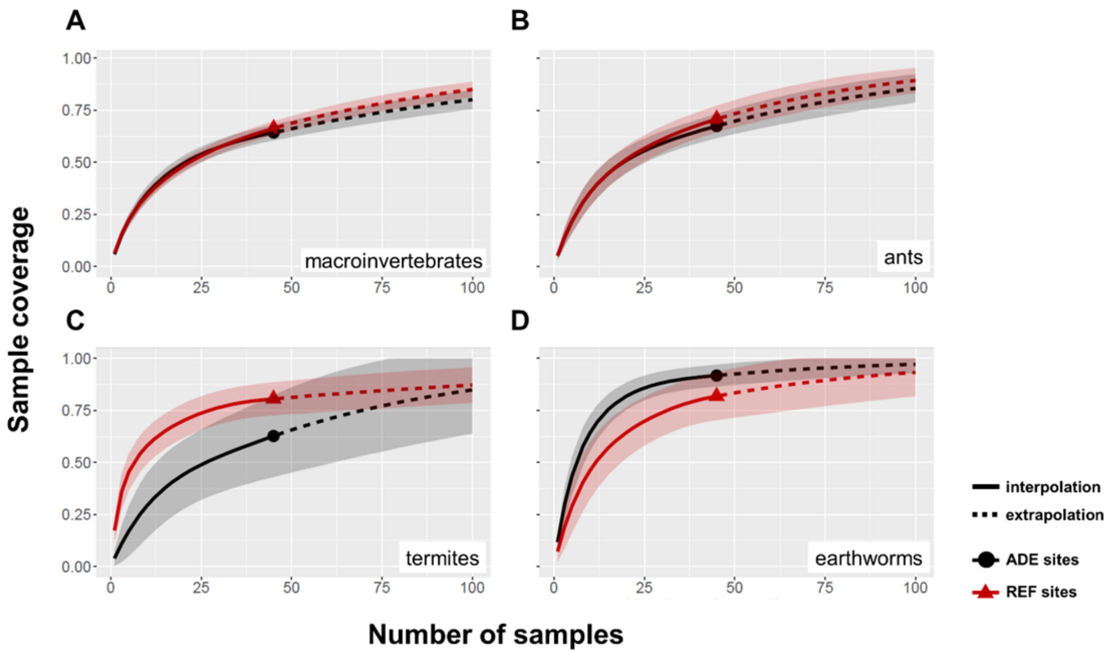

**Supplementary Figure 6. Sampling effort coverage**, showing diversity collected depending on sampling intensity (number of samples) in ADEs and REF soils for: (**a**) All soil macroinvertebrates, (**b**) ants, (**c**) termites and (**d**) earthworms. Data correspond to invertebrates collected using soil monoliths, over all sites and land use systems. Dark grey and red areas represent 95% confidence intervals. REF: Reference soil, ADE: Amazonian Dark Earth soil

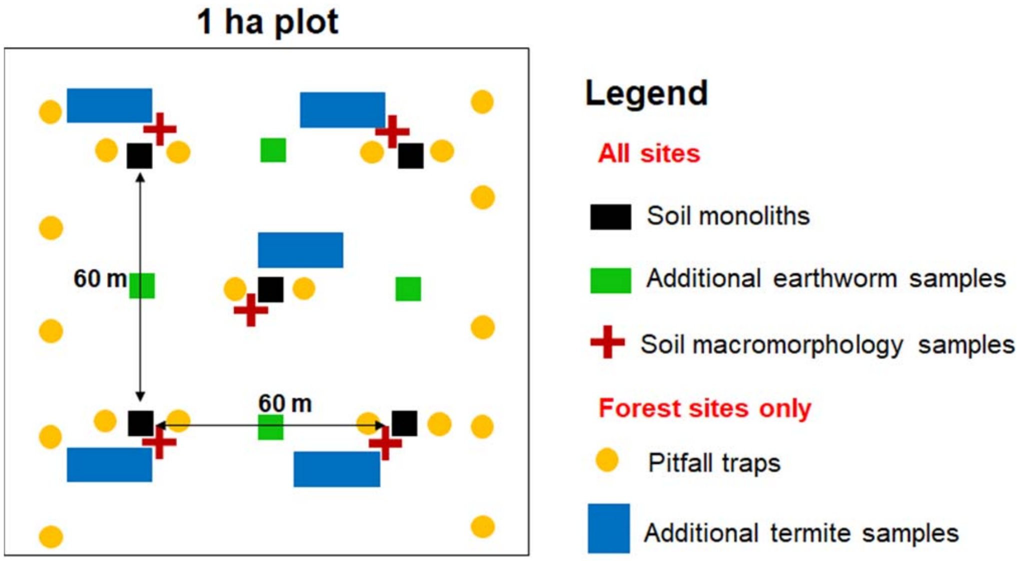

**Supplementary Figure 7. Scheme used for soil and fauna sampling** for each plot in each land use system. Distribution of the different samples in the 1 ha plot at all sites, showing the types of samples taken: monoliths for soil fertility and total soil macrofauna, soil macromorphology samples and additional samples for earthworms, and forest-only samples for termites (2 x 5 m plots) and ants (pitfall traps).
